# Telomere to telomere sequence of model *Aspergillus fumigatus* genomes

**DOI:** 10.1101/2022.03.26.485923

**Authors:** Paul Bowyer, Andrew Currin, Daniela Delneri, Marcin G. Fraczek

## Abstract

The pathogenic fungus *Aspergillus fumigatus* is a major etiological agent of fungal invasive and chronic diseases affecting tens of millions of individuals worldwide. A high-quality reference genome is a fundamental resource to study its biology, pathogenicity and virulence as well as to discover better and more effective treatments against diseases caused by this fungus. Here, we used PacBio Single Molecule Real-Time (SMRT) and Oxford Nanopore sequencing for *de novo* genome assembly of two laboratory reference strains of *A. fumigatus,*CEA10 and A1160. We generated full length chromosome assemblies and a comprehensive telomere to telomere coverage for these two strains including ribosomal repeats and the sequences of centromeres, which we discovered to be composed of long transposon elements.

## INTRODUCTION

*Aspergillus fumigatus* causes over 11 million allergic and over 3 million chronic and invasive lung infections annually, representing a significant complication of profound immunosuppression, chronic obstructive pulmonary disease (COPD), severe viral respiratory infections (such as influenza or Covid-19) and many other pre-existing conditions (1–4). Mortality rates with effective treatment for invasive disease remain ~50% (5) and >80% for individuals infected with drug resistant isolates (6). The availability of genome sequence has underpinned many of the rapid advances in our understanding of this organism in recent years.

The first *A. fumigatus* genome sequence was published in 2005 (7) for a clinical isolate Af293, followed by the A1163 strain published in 2008 (8). These two reference genome sequences have been crucial to study the biology and pathogenicity of this fungus. However, due to the technological capabilities at the time, the original reference sequences are not complete and contain several sequence blocks of insertions and duplications, some sequences are absent (deletions) or unknown nucleotides (NNN) are present. Moreover, these sequences lack coverage of centromeres, the accurate sequence for the ribosomal repeats, and a comprehensive annotation of chromosomal rearrangements such as translocations and inversions. In particular, A1163 was not sequenced at sufficient depth to perform a full assembly and remains as poorly organised contigs and scaffolds (8). A1163 or strains derived from its parental isolate CEA10 (9, 10) have become standard in laboratory experiments because of their robust pathogenicity and growth. For example, the CEA10 descendant isolate A1160, recently renamed to MFIG001 (10), is a laboratory isolate in continuous use nowadays, first mutated to uridine auxotrophy (*pyrG^−^*) to form CEA17 (11) and subsequently used to construct the *pyrG*+ *ku80* knockout strain (12). A1160 is currently being used as a host strain for a whole genome knockout project (13, 14) and forms the basis of many virulence, transcriptome and other experiments (15). Therefore, there is an urgent requirement to revise the original genome sequences and provide comprehensive genome assemblies of the most exploited *A. fumigatus* strains A1160 and CEA10 using the current long-read sequencing technology.

Recent advances in the third-generation sequencing technologies, such as Pacific Biosciences (PacBio) and Oxford Nanopore, allow longer reads and more accurate assembly of genomic sequences. They have been used to provide complete and accurate genome assemblies of a wide range of organisms, including human, plants and animals as well as fungal pathogens such as *Magnaporthe oryzae* and *A. awamori* (16–18). Due to the pathogenic nature of *A. fumigatus*, with large numbers of patients suffering from aspergillosis worldwide as well as increasing numbers of fungal studies, there is an urgent requirement for the assembly of high-quality reference genomes of commonly used *A. fumigatus* isolates. Therefore, here we present the complete high quality *de novo* telomere to telomere sequences of CEA10 and its descendant isolate A1160, revealing centromere structure, ribosomal repeat sequence and chromosomal organisation. As previously predicted (8), CEA10 shows chromosomal rearrangements when compared to Af293 and there is evidence of a small number of mutations affecting gene function that have accrued in the last ~30 years since isolation of CEA10, and the creation of A1160 in the laboratory. The sequences analysed in this study will be publicly available for the scientific community and will greatly contribute to the future research on this fungus.

## RESULTS AND DISCUSSION

### Sequencing and *de novo* genome assembly

The complete genome sequence of two *A. fumigatus* laboratory reference strains, A1160 and CEA10 was carried out using the long read *de novo* PacBio and Oxford Nanopore third generation sequencing technologies. The data acquired allowed us to greatly improve the quality of the genome assembly compared to the original reference sequences of Af293 and A1163 (7, 8) and expand the genomic resources for this pathogen. Specifically, missing gaps were filled and additional genomic information on ribosomal repeats and centromere composition was added. Interestingly, we found that the centromeres of *A. fumigatus* encompass long stretches of DNA and are composed of transposons. Moreover, the comparative analysis of A1160 and CEA10 vs Af293 revealed several chromosomal rearrangements, the largest of which being detected on chromosomes 1 and 6.

The PacBio and Oxford Nanopore sequencing generated sufficient data to allow high quality genome assemblies of the expected >29 Mb size (7). Both strains were assembled in 10 contigs with 233x and 183x coverage for A1160 and CEA10 genomes, respectively, using the PacBio assembly algorithms (Supplementary Dataset 1). GC content for both strains was ~49.5%. For the Oxford Nanopore sequencing, the same genomic DNA was used unsheared which provided longer raw data reads with N50 of 20 kB. The mean coverage for the Oxford Nanopore assembly using Canu 1.9 was 39x for both strains with 23 and 19 contigs for A1160 and CEA10, respectively (Supplementary Dataset 1).

Our data show that the genomes of A1160 and CEA10 are almost identical in sequence besides a small number of SNP variations (96) in several genes (Supplementary Dataset 2). The most evident changes in the SNPs are observed on chromosome 8, for which we also observed several insertions and deletions (INDELs) of nucleotides, leading to frame shift. There is a total of 34 INDELs between these 2 strains. For the strain A1160 the telomere on chromosome 6 could not be completely assembled due to chromosomal rearrangements.

Ribosomal sequence was extracted from the raw data using grep to capture reads known to contain *A. fumigatus* ribosomal sequences. For Oxford Nanopore data, assembled repeat regions were obtained as assembled contigs. The core assembly indicated only a single 28S repeat and this is likely due to mis-assembly of the repeat units. As the number of repeats is not clearly distinguishable, the 28S segment was left as a marker for the region on chromosome 4. Assembled ribosomal repeats and the minimal repeat unit are provided in the .fasta files (Supplementary Dataset 3 and 4).

The mitochondrial sequences of both species were also analysed, and we found that our assembled data are consistent with previously published sequences for A1160 and Af293 (19).

### The new genome assembly unravel previously undetected gene sequences and chromosomal rearrangements

The original sequence of Af293 was created in 2005 using the whole genome random sequencing method (7). Although, it still provides crucial sequencing data, it does not include centromeres or chromosomal rearrangements. In table 1 we summarise the predicted sizes of chromosomes and genes from our PacBio analysis for A1160 and CEA10 and compare them to the sizes present in the database for Af293. Two different pipelines were used in this analysis revealing no major differences in the sizes between the previously generated reference sequences and our newly assembled genomes. As previously shown (7), the genome o *A. fumigatus* is arranged in 8 chromosomes of a total of approximately 29.2 Mb. However, compared to the previously sequenced Af293, we found that A1160 and CEA10 have approximately 300 more genes (Table 1), which are the result of more accurate assembly.

**TABLE 1.** Assembly statistics for assembled chromosomes. Chromosome sizes in base pairs (bp). Genes were predicted for each chromosome either by using a GenemarkEP+/Braker pipeline (GMEP/Braker) or via transfer of annotations from the Af293 FungiDB database using exonerate (FDB).

| Chromosome<br>(bp) | A1160 |  |  | CEA10 |  |  | A1163 |  |
| --- | --- | --- | --- | --- | --- | --- | --- | --- |
|  | Size (bp) | Gene Count |  | Size (bp) | Gene Count |  | Size (bp) | Gene Count |
|  |  | GMEP/Braker | FDB |  | GMEP/Braker | FDB |  |  |
| 1 | 4910149 | 1684 | 1671 | 4910137 | 1671 | 1671 | 4918979 | 1612 |
| 2 | 4786411 | 1617 | 1634 | 4786983 | 1607 | 1635 | 4844472 | 1624 |
| 3 | 4271837 | 1479 | 1453 | 4270431 | 1477 | 1452 | 4079167 | 1362 |
| 4 | 3812114 | 1288 | 1290 | 3819196 | 1285 | 1289 | 3923705 | 1224 |
| 5 | 3799872 | 1339 | 1345 | 3799900 | 1330 | 1345 | 3948441 | 1351 |
| 6 | 3848548 | 1332 | 1282 | 3849020 | 1330 | 1281 | 3778736 | 1227 |
| 7 | 1884753 | 617 | 608 | 1884962 | 614 | 608 | 2058334 | 621 |
| 8 | 1944132 | 641 | 631 | 1949190 | 642 | 636 | 1833124 | 609 |
| Total | 29257816 | 9997 | 9914 | 29269813 | 9956 | 9917 | 29384958 | 9630 |

Protein coding gene transcripts, short ncRNAs, tRNAs as well as transposons were annotated based on our *de novo* analysis and the data from FungiDB (Fig. 1; Supplementary Dataset 5 and 6). When determining centromere localisation, we observed that transposable elements, besides being scattered throughout the whole genome as predicted, they were also localised in the centromeres of all 8 chromosomes, forming the majority of centromeric sequences. Although, it was previously predicted that centromeres of filamentous fungi may be composed of transposons (20), our study is the first to confirm that the centromeres of *A. fumigatus* chromosomes are enriched with transposable elements. An example of a detailed chromosomal annotation is presented in Fig. 2.

**Figure 1.**
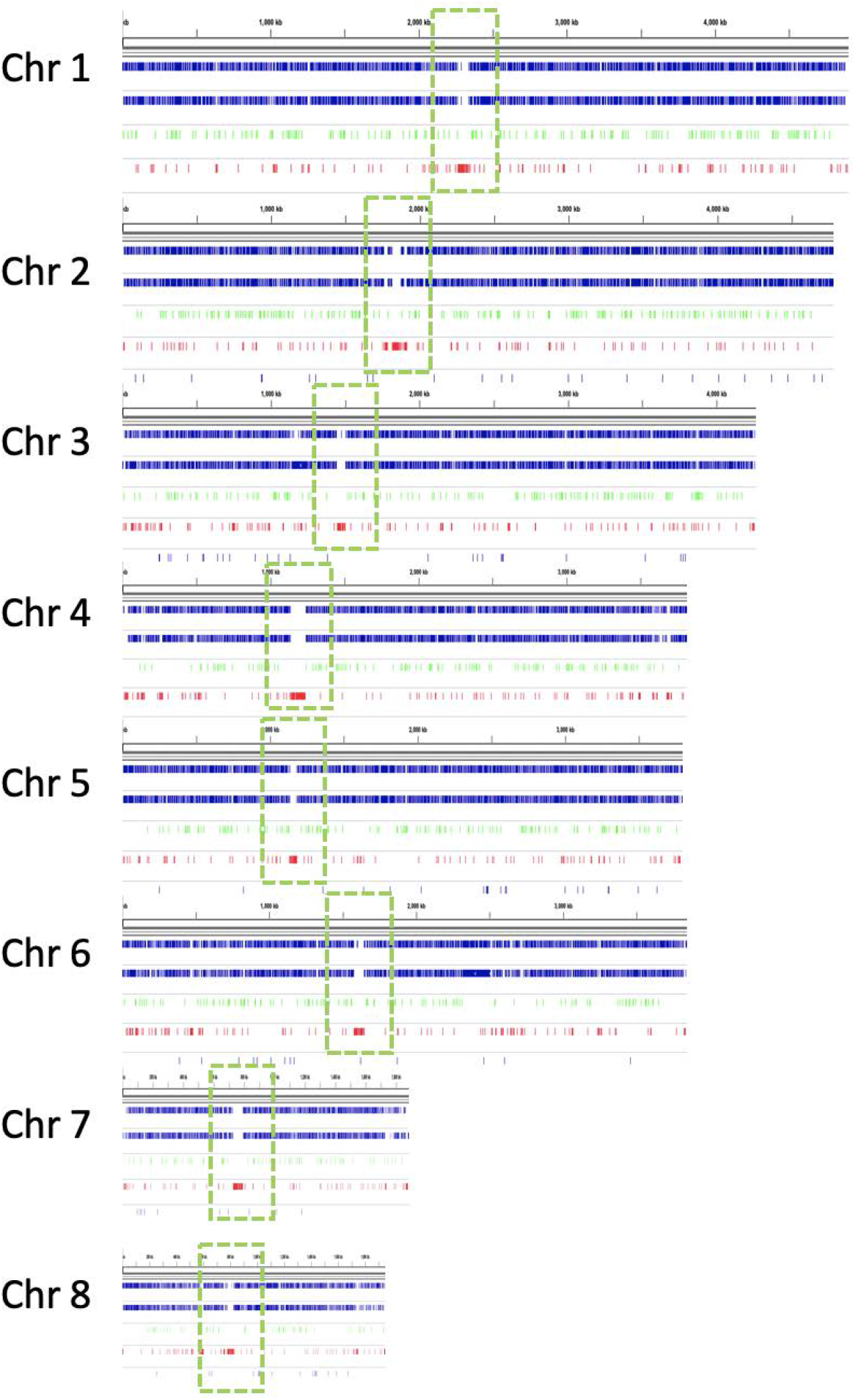
Annotation of A1160 genome assembled in eight chromosomes. Putative centromeres are indicated by green boxes and genes are shown in blue. Genomes are annotated in a cursory manner using exonerate mapping of transcripts and proteins from Fungi DB and *de novo* gene prediction using Genemark EP+ and Braker1 and Braker2. Transposons are mapped using exonerate (Transposons) and their positions is shown in red. Short RNAs such as tRNA are also mapped in light green. Random PCR RNAseq data from NCBI was merged and mapped to genomes using HiSat2.

**Figure 2.**
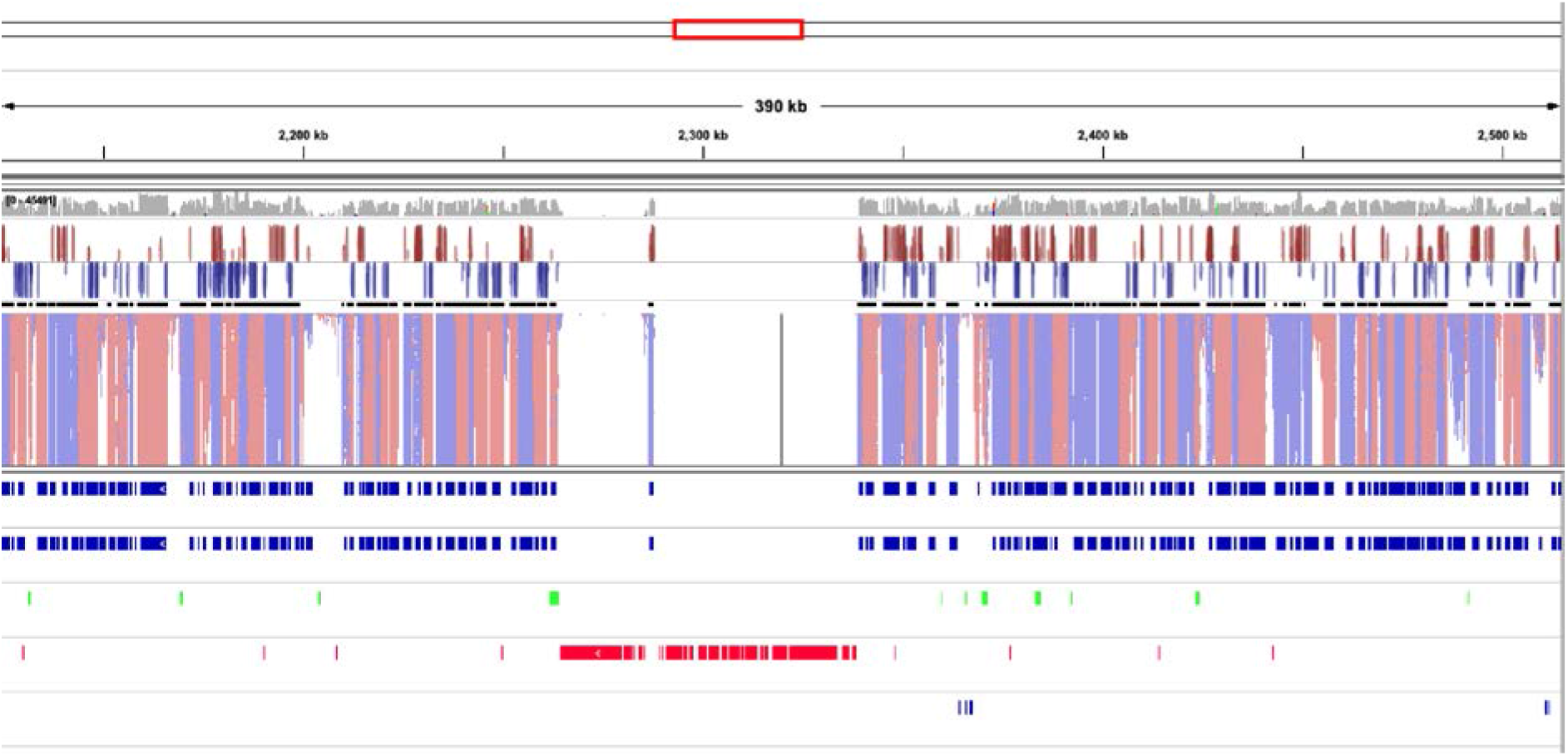
Annotation of CEA10 genome showing a detail of chromosome with a putative centromere. Genes are shown as blue boxes. Genomes are annotated in a cursory manner using exonerate mapping of transcripts and proteins from (Fungi DB) and *de novo* gene prediction using Genemark EP+ and Braker1 and Braker2. Transposons are mapped using exonerate (Transposons) with positions shown in red. Long non-coding RNA and short RNAs such as tRNA are also mapped and shown in light green. Random PCR RNAseq data from NCBI was merged and mapped to genomes using HiSat2.

Our sequencing data also confirmed the localisation of the native *ku80* gene deletion in CEA10 (9) as well as the replacement of this gene in A1160 with *pyrG^+^* on chromosome 2 (12) (Fig. 3).

**Figure 3.**
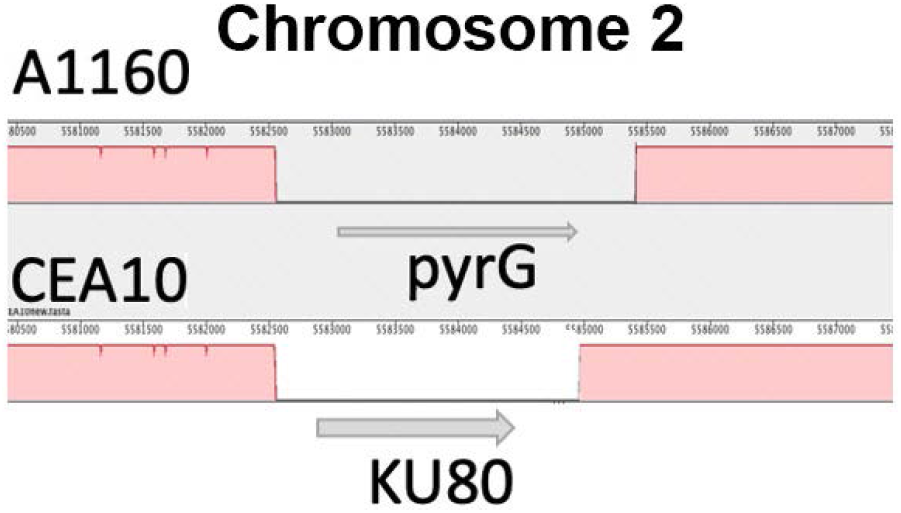
Zoom-in of the chromosome 2 map showing gene replacement of *ku80* in the A1160 strain with reference to the parental CEA10 isolate.

The comparison between the genomes of the reference strain Af293 and sequenced CEA10/A1160 revealed a number of chromosomal rearrangements (Fig. 4A and B). The largest rearrangements are between the ends of chromosomes 1 and 6 (a situation previously suggested in the original A1163 sequencing (8)). Chromosomal rearrangements and chromosomal breakpoint usage have been proposed to play a significant role in evolution that lead to environmental adaptation and these events have been previously observed in filamentous fungi (21–23). As both A1160 and CEA10 strains have been widely used for >20 years, it is expected that they might have accrued mutations and chromosomal rearrangements.

**Figure 4.**
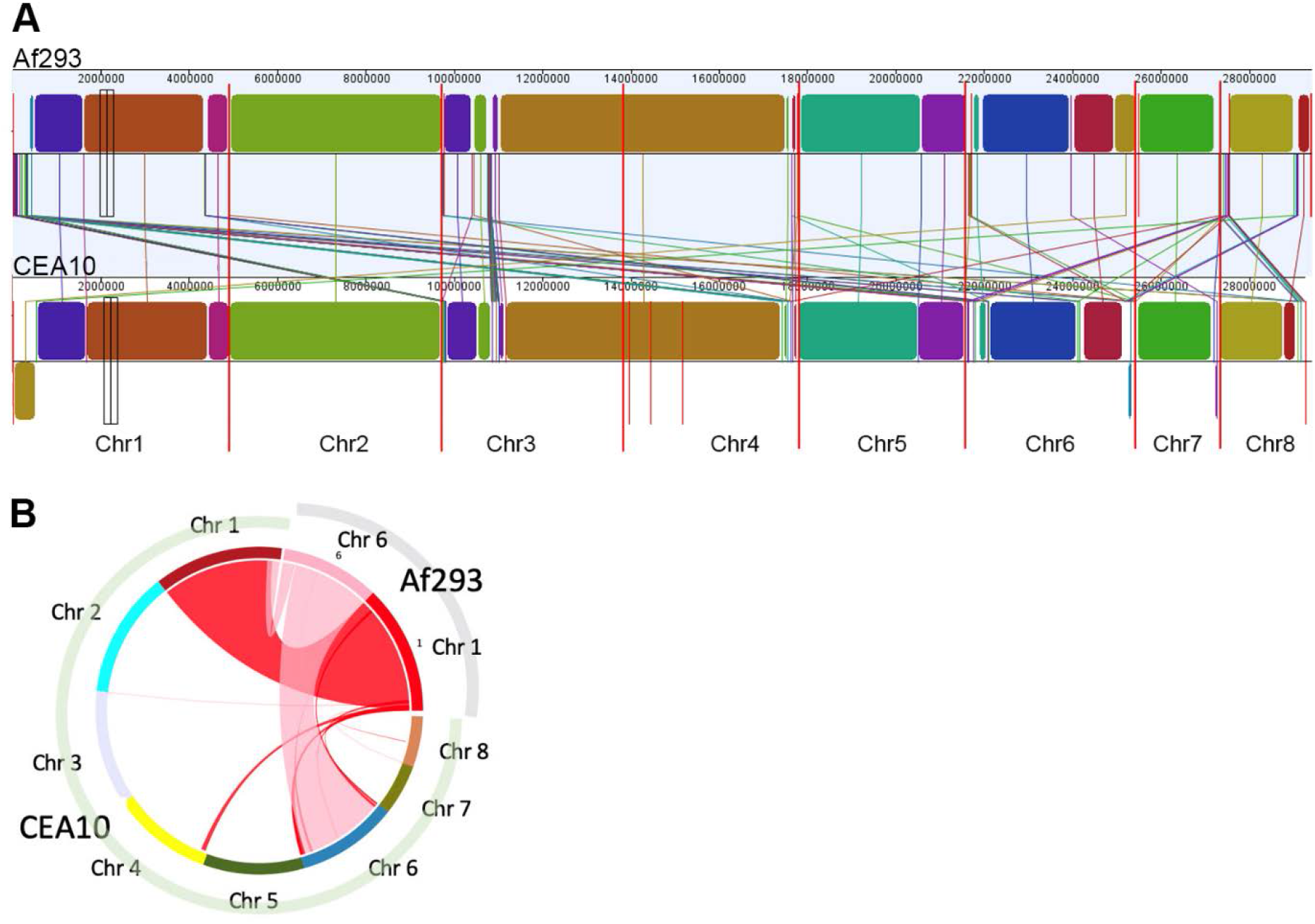
Chromosomal rearrangements observed between Af293 and CEA10. (A) Chromosomes are marked along the bottom of the panel. Syntenic blocks are shown as different coloured boxes for both strains with identical colours indicating synteny. Connecting lines are shown to indicate chromosome rearrangement. (B) Circular plot showing detailed chromosome rearrangements between chromosome 1 and 6 in both Af293 and CEA10 strains.

## CONCLUSIONS

The availability of comprehensive genome sequence of *A. fumigatus* strains is crucial to understand the biology, pathogenicity and virulence of this fungus. Moreover, quality genome sequences are proving to be a powerful method for discovering mechanisms of drug resistance and may lead to more efficient patient treatment and their recovery. Here, we provide the comprehensive, telomere to telomere genome sequences of two widely used isolates of *A. fumigatus,* A1160 and CEA10, to fill in the gaps in the sequences of the original reference strains, Af293 and A1163. Our data shows significant improvement in sequence quality and organisation of chromosomes, revealing centromere structures and ribosomal repeats. The assembled sequences in this study are of great interest to the scientific communities that lead research into better treatment and diagnostics of fungal diseases.

## MATERIALS AND METHODS

### Strains and genomic DNA preparation

Two strains of *A. fumigatus,* CEA10 and A1160 (10, 12) were used in this study. Fungal spores were used to extract high quality genomic DNA following a previously described CTAB method (12) with few modifications that greatly improved the quality and purity of extracted DNA. Briefly, both isolates were grown on SAB agar media in tissue culture flasks to minimise cross-contamination and spores were harvested in PBS/Tween20 and transferred to 2 ml screw top tubes containing 425–600 mm washed glass beads (filled to the 300 mL mark; ~50 mg) (Merck). Spores were centrifuged at max speed for 2 min using a benchtop centrifuge and the supernatant was removed. 1 mL of CTAB extraction buffer (2% CTAB, 100 mM Tris, 1.4 M NaCl and 10 mM EDTA, pH 8.0) was added and the tubes and they were vortexed at max speed for 10 minutes. Subsequently, the tubes were incubated for 10 min at 65°C. Then, the above vortexing and heating process was repeated, and tubes were centrifuged at max speed for 2 minutes. The supernatant was transferred to new 2 ml tubes and an equal volume of chloroform was added. Tubes were mixed by inversion and centrifuged for 3 minutes at max speed. The aqueous phase was transferred to new 1.5 mL tubes and DNA was precipitated by addition of 0.6 volumes of isopropyl alcohol. Following centrifugation for 2 minutes at max speed, the supernatant was decanted, and the pellet was washed with 0.5 mL absolute ethanol. The pellet was briefly air-dried and resuspended in 200 μL of dH_2_O. Subsequently, 2 μl of 100 mg/mL RNase A (Qiagen) was added and the tubes were incubated at 37°C for 15 minutes. Then, 1 mL of buffer PB or PM (Qiagen), containing a high concentration of guanidine hydrochloride and isopropanol was added and mixed by pipetting. The solution was transferred onto silica based blue columns (NBS biologicals) and centrifuged for 30 seconds at max speed. Then, 700 μL of buffer PE (Qiagen) was added onto the column and centrifuged as above followed by additional spinning for 1 minute at max speed. The DNA was eluted in 100 μL of dH2O and the quality of the DNA was assessed on a 1% agarose gel, as well as using a nanodrop (Thermofisher Scientific) and a Qubit 4 Fluorometer (Thermofisher Scientific).

### Library preparation for long read next generation sequencing

For PacBio sequencing, genomic DNA was adjusted to 10 ng/μL in 150 μL volume and sheared to approximately 10 kb fragments using g-TUBES (Covaris) following the manufacturers’ instructions. The size of fragments and quality of the DNA was verified using a Fragment Analyzer (Agilent) and the DNF-930 protocol. Samples were prepared for sequencing following the Express Template Prep Kit 2.0 protocol, with multiplexing using the Barcoded Overhang Adapter kit 8A (both Pacific Biosciences). DNA libraries were sequenced using the SMRT Cell 1M chips on the Pacific Biosciences Sequel system with 10 hour data acquisition time.

For Oxford nanopore sequencing, 1 μg of the same DNA samples (not sheared) were prepared for sequencing using the SQK-LSK109 Ligation sequencing kit and Flongle sequencing expansion kit, following the manufacturer’s instructions. Each strain was sequenced using a MinION Flongle flow cell with 24 hours data acquisition time.

Illumina paired end reads were also obtained using a HiSEQ2500 in order to provide better sequence coverage.

### Genome assembly

Demultiplexing and *de novo* assembly was performed using the Pacific Biosciences algorithms within the SMRT Link 8.0 software package. For *de novo* assembly the Hierarchical Genome Assembly Process (HGAP4) was used, with 30x seed coverage specified for each assembly with specified genome length of 29 Mb (all other parameters were unchanged). Assembly polishing and resequencing was performed using the Resequencing algorithm in SMRT Link 8.0.

For Oxford Nanopore data, base calling was performed using Guppy (Oxford Nanopore) and *de novo* assembly was performed using Canu 1.9 (24), with specified genome length of 29 Mb.

For CEA10 PacBio and Nanopore sequence assemblies were then polished using 3 rounds of PILON (25) with 2 paired end Illumina 2×150 fastqs to give the final CEA10 sequence.

### Annotation

Genomes were subjected to a cursory annotation using a Genemark EP+ pipeline (26) guided by Prothint 2.5.0 using orthodb version 10.1 as previously described (27). Additionally, Augustus, BRAKER1 and 2 annotations were performed according to the software defaults for *A. fumigatus* and fungi, respectively (28). Finally, an existing curated annotation for A1163 was mapped to the A1160 and CEA10 genomes using Exonerate (29). Transposon sequences were collected for *A. fumigatus* from NCBI searches and further mapped onto the genome sequences using Exonerate. Transcript data from NCBI SRA archive was used to guide annotation and to generate a list of potential transcribed regions which were then tested for the presence of ORFs, ORFs matching known proteins in the UniRef90 dataset or ORFs with matching PFAM domains using TransDecoder (https://github.com/TransDecoder/TransDecoder). Annotations and cognate assembled genome sequences are available in Supplementary Dataset 5 and 6.

## Supporting information

Supplementary Dataset 1 - Assembly metrics

Supplementary Dataset 2 - SNPs-INDEL

Supplementary Dataset 4 - CEA10

Supplementary Dataset 6 - CEA10

## Acknowledgements

The authors would like to thank Michael Bromley for sourcing the CEA10 strain. MF has been supported by Wellcome Trust, under the grant number 208396/Z/17/Z.

---

**Supplementary Dataset 1** - Assembly metrics for Pacific Biosciences and Oxford nanopore data.

**Supplementary Dataset 2** - Variation between CEA10 and A1160. 130 variants with predicted high, moderate or low impact on gene function are listed from 395 supported variants in the comparison.

**Supplementary Dataset 3** – Assembled A1160 sequence (.fasta file)

**Supplementary Dataset 4** - Assembled CEA10 sequence (.fasta file)

**Supplementary Dataset 5** – Annotations and cognate assembled genome sequences of A1160 (.gff file)

**Supplementary Dataset 6** – Annotations and cognate assembled genome sequences of CEA10 (.gff file)

